# Visual intensity ratio modulates operant learning responses in larval zebrafish

**DOI:** 10.1101/401000

**Authors:** Wenbin Yang, Yutong Meng, Danyang Li, Quan Wen

## Abstract

Larval zebrafish is a promising vertebrate model for understanding neural mechanisms underlying learning and memory. Here, we report on a high-throughput operant learning system for zebrafish larvae and demonstrate that lower visual intensity ratio of the conditioned stimulus to the background can enhance learning ability, highlighted by several behavioral metrics. We further characterize the learning curves as well as memory extinction for each conditioned pattern. Finally, we show how this learning process developed from 7 days old to 10 days old zebrafish.

**Highlights:**

- Conditioned visual patterns with lower intensity ratio to the background elicited stronger operant learning responses
- Memory extinction was modulated by the visual intensity ratio of the conditioned stimulus to the background
- A high-throughput automated system for acquiring and analyzing behavioral data

## 1 Introduction

In operant conditioning, an animal learns, through trial and error, to correlate its behavioral responses with the consequences. This form of associative learning has been intensively studied in mammals (Freund and Walker, 1972; Ishikawa et al., 2014), but the biological learning rules, as well as their implementation by the brain circuit, remains elusive. To make progress, it would be illuminating to measure neural activities of defined cell types at the whole-brain scale during the entire learning process. Larval zebrafish is a promising vertebrate model for this purpose: the transparency and the relatively small brain is a great compromise between system complexity and simplicity. Recently, it has become possible to perform the whole-brain imaging of calcium activities in freely behaving larval zebrafish (Cong et al., 2017; Kim et al., 2017). Whereas fish are well-established animal models to study learning and memory (Agranoff and Davis, 1968; Davis and Agranoff, 1966), few associative learning paradigms have been developed for zebrafish larvae. Li (Li, 2012) reported operant learning in head-fixed larvae with aversive heat stimulus; Valente and colleagues (Valente et al., 2012) showed that one-week larvae were unable to perform an operant learning paradigm, in which fish must learn to swim to the other half of an arena to avoid electroshocks. Other reports demonstrated that larval zebrafish could also learn classical conditioning: they could associate the conditioned stimulus (CS), a moving spot, with the unconditioned stimulus (US), a touch of the body (Aizenberg and Schuman, 2011). Social reward, such as visual access to conspecifics, could also be paired with a distinct visual environment during classical conditioning in larval zebrafish (Hinz et al., 2013).

Zebrafish have sophisticated vision: they can discriminate size, color, intensity and object motion with ease. Spatial and non-spatial visual learning tasks have been well studied in adult zebrafish (Arthur and Levin, 2001). However, much less is known about how visual properties would modulate the learning process in larval zebrafish. Here, we reported a modified operant conditioning paradigm (Valente et al., 2012) in freely swimming larval zebrafish, a system that combines a high-throughput automated training process and a toolkit for post-data analysis and storage. We used our new paradigm to investigate how visual intensity ratio modulated the operant learning responses in larvae, characterized by both positional and turning metrics. We also quantified the learning curves and memory extinction for individuals.

## 2 Material and Methods

### 2.1 Ethical statement of animals-using

Handling and care of all animals were conducted in strict accordance with the guidelines and regulations set forth by University of Science and Technology of China (USTC) Animal Resources Center, and University Animal Care and Use Committee. Both raising and training protocols were approved by the Committee on the Ethics of Animal Experiments of the USTC (permit number: USTCACUC1103013).

### 2.2 Animals and raising

Zebrafish (*Danio rerio*) of the genotype huc:h2b-gcamp6f were used in all experiments. All tested fish were from 7 to 10 dpf (day past fertilization) larvae. They were bred, raised and housed in the same environment. Fish were fed two times per day from 6 dpf with paramecium in the morning (8-9 A.M.) and evening (6-7 P.M.) until used in the experiments. Water was replaced with E2 medium (Cunliffe, 2003) in the morning (8-9 A.M.) and evening (6-7 P.M.). Water temperature was maintained at 28.5 °C. Illumination was provided by fluorescent light tubes from the ceiling with lights turned on at 08:00 A.M. and off at 10:00 P.M.

### 2.3 Experimental Setup

The behavioral system with custom software suites and supported hardware were built to achieve an end-to-end high-throughput experimental workflow. (Figure 1A)

**Figure 1.**
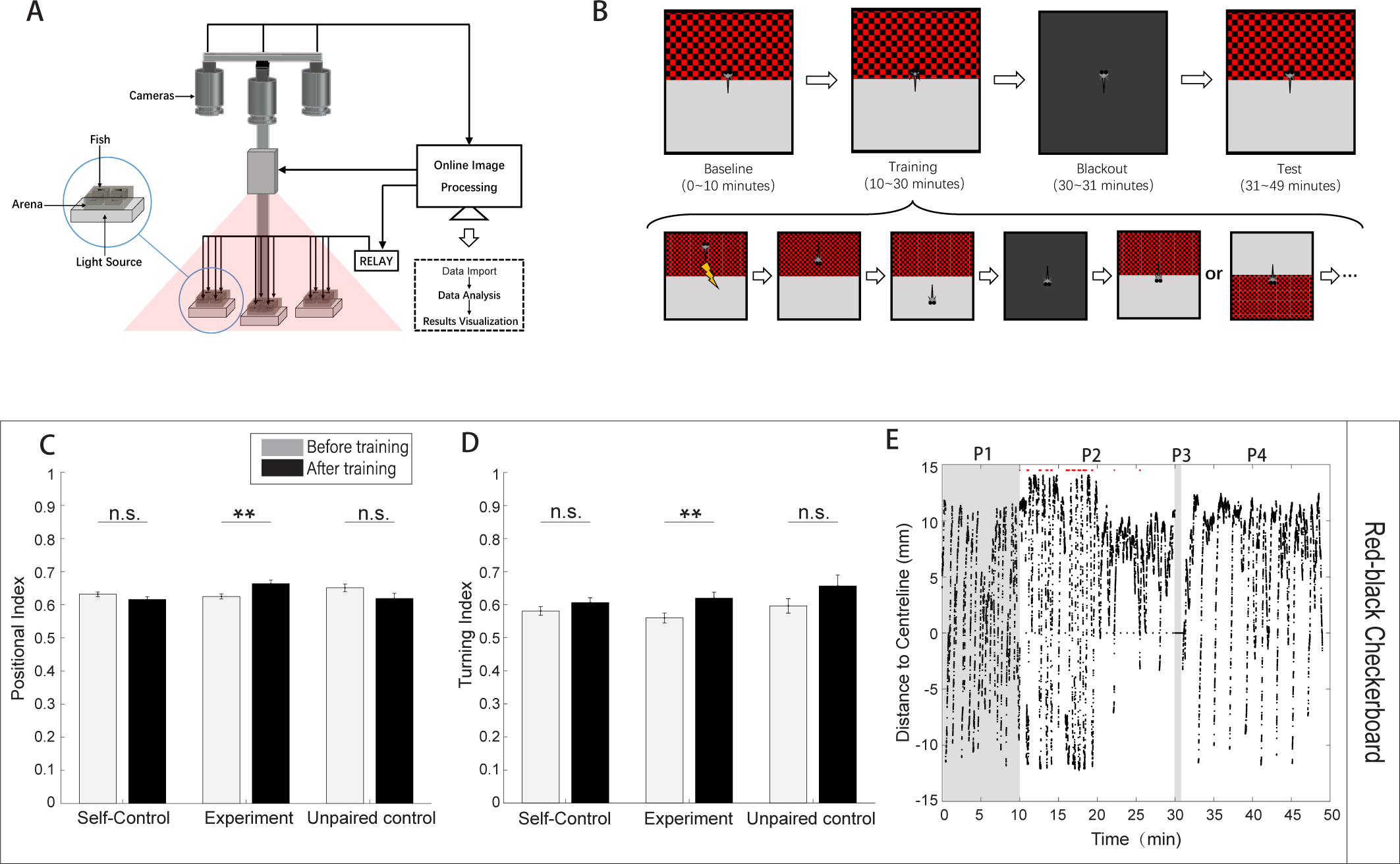
Larval zebrafish show significant learning responses in the operant learning task. (A) Schematics of the behavioral system. Each arena holds one fish. Cameras, projector, relay are controlled by the custom software BLITZ. Software ABLITZER imports the BLITZ-produced behavioral data, analyzes them and visualizes the results. (B) Operant learning paradigm. The procedure (top) and the detailed protocol in the training stage (bottom). (C) Larval zebrafish showed significant learning responses, quantified by the positional index, after operational conditioning. Each bar pair shows the positional indices before (light bar) and after (black bar) the training (p = 0.0843, p = 0.0021, and p = 0.1260 from left to right, t-test). (D) Larval zebrafish showed significant learning responses, quantified by the turning index, after operational conditioning (p = 0.1836, p = 0.0099, and p = 0.0628, t-test). (E) A representative behavioral trace. A typical learner’s relative position to the midline during an operant learning experiment (CS zone: red-black checkerboard, non-CS zone: pure gray pattern). A positive distance suggests fish in the non-CS zone (also see Methods). Each red dot represents the delivery of one electroshock.

#### 2.3.1 Hardware

Zebrafish swam freely in custom-built acrylic containers with transparent bottoms. Each container was divided into four arenas separated by opaque walls. The arena’s size is 3 cm × 3 cm × 1 cm, with water filled (Fig. S2.a). Each arena held one fish. Three CMOS cameras (Basler aca2000-165umNIR, Germany) with adjustable lens (Canon, Model EF-S 18-55mm f/3.5-5.6 IS II, Japan) simultaneously captured swimming behavior at ten frames per second. Three infrared LED light sources (Kemai Vision, China, model HF-FX90, wavelength 940 nm) illuminated each container from below. A 700 nm long-pass filter (Thorlabs FEL0700, US.) was positioned in front of each camera to block visible light to facilitate online imaging processing with custom software BLITZ. Visual stimuli were presented by a projector from the top over all three containers (PIQS Projector S1, 14.6 × 7.85 × 1.75 cm, 854 × 480 pixels). Electroshocks (100 ms, 9 Volt/3 cm) were delivered via two platinum filaments, one on each side of the arena. Shock delivery at each arena was controlled by custom software BLITZ via a 16-channels relay (HongFa JQC-3FF, China). Room temperature was controlled by an air-conditioner at 27 °C.

#### 2.3.2 Software Suites

Custom C++ software BLITZ (Behavioral Learning In The Zebrafish, inheriting the coding style from MindControl (Leifer et al., 2011)) with Microsoft Visual Studio 2017 processed three video streams in parallel to get real-time head, center, tail positions and heading angle by using the Pylon library (Basler AG, Germany) and the open source computer vision library (OpenCV) (Bradski, 2000). The program also rendered visual pattern and programmable electroshocks delivery based on the timeline and real-time fish motion parameters. All necessary experimental information (e.g., experiment start time, visual pattern index, shocks delivery information, and fish motion parameters) were recorded in YAML files. Raw videos were recorded.

The BLITZ software is available at https://github.com/Wenlab/BLITZ.

Another custom MATLAB (The MathWorks, Inc.) software ABLITZER (the Analyzer of BLITZ Results) was used to import YAML files, to visualize data, as well as to perform the behavioral and statistical analysis.

The ABLITZER software is available at https://github.com/Wenlab/ABLITZER.

### 2.4 Experimental Procedure

Fish were fed at least an hour before using in the experiment. Fish were placed via a Pasteur pipette (Nest, US) from the raising tank to the experimental arenas. The behavioral experiment would not run until fish started moving around to avoid startle responses to novel stimuli. Fish in the paired-group were trained first with the self-control protocol (see below), then with the operant learning protocol. Fish in the unpaired-group were trained first with the self-control protocol, then with the unpaired operant learning protocol (see below).

Fish used in the paired-group and unpaired-group were all naive fish before the experiment.

#### 2.4.1 Operant learning protocol

This operant learning protocol was modified from Valente’s learning paradigm (Valente et al., 2012). Here, fish would experience four different phases in order: baseline phase, training phase, blackout phase and test phase. (Figure 2B)

**Figure 2.**
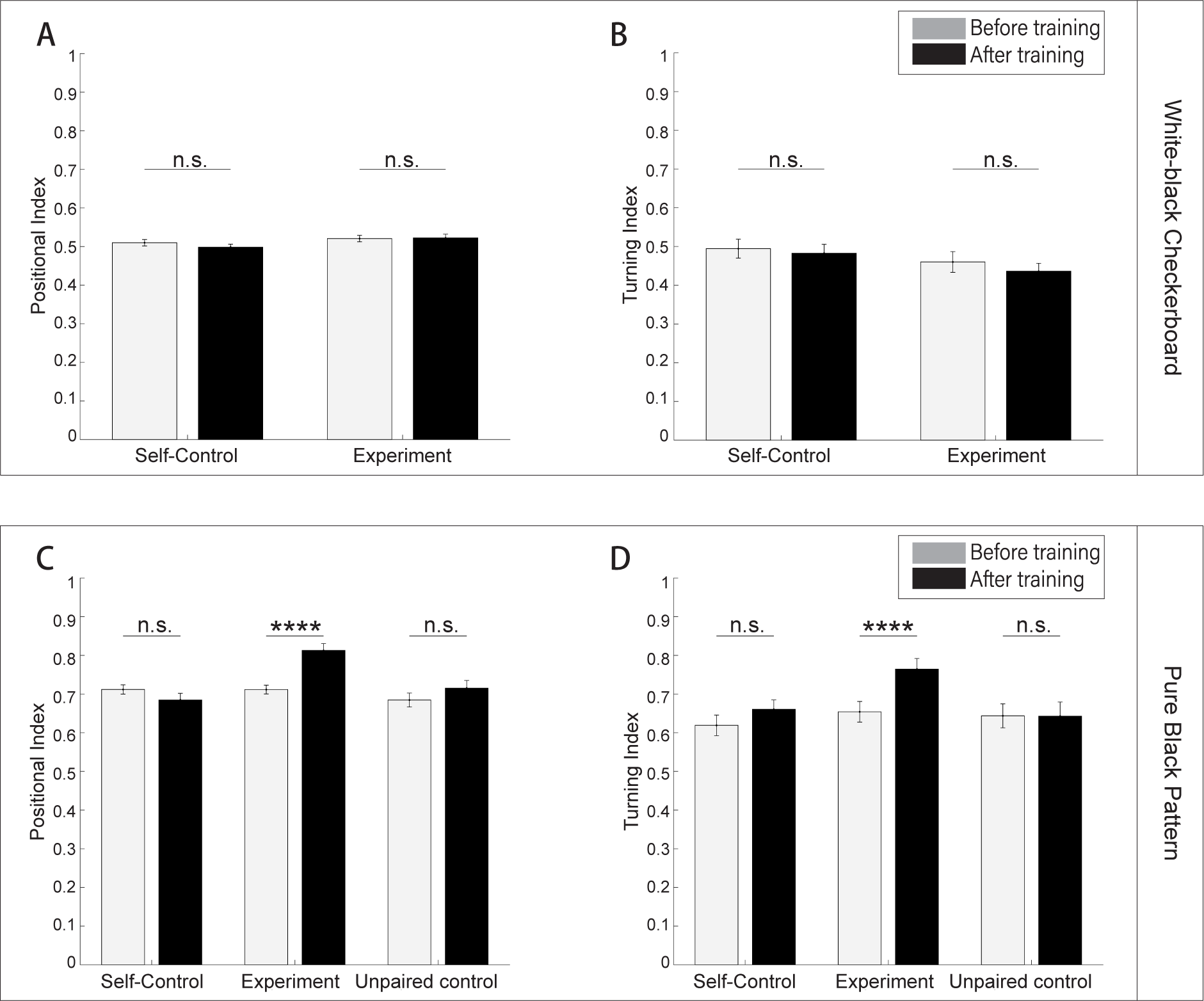
Visual intensity ratios modulate operant learning responses in larval zebrafish. (A) Analysis of the positional index suggested that fish did not show significant learning responses (CS zone was white-black checkerboard). t-test, p = 0.8832 for the experiment group, p = 0.2493 for the self-control group. There is no unpaired-control group because no significant learning responses were found in the experiment group. (B) Analysis of the turning index suggested that fish did not show significant learning responses (CS zone was white-black checkerboard). t-test, p = 0.3750 for the experiment group, p = 0.7089 for the self-control group. There is no unpaired-control group because no significant learning responses were found in the experiment group. (C) Analysis of the positional index suggested that fish showed significant learning responses. (CS zone was black pattern; p = 0.2018, p < 0.0001, p = 0.1923, from left to right respectively, t-test.) (D) Analysis of the turning index suggested that fish showed significant learning responses. (CS zone was black pattern; p = 0.2811, p = 0.0057, and p = 0.9837, from left to right respectively, t-test.)

First, in the 10 minutes baseline phase, the visual pattern beneath each arena would flip between the CS at the top (Figure S1A, C or E) and the CS at the bottom (Figure S1B, D or F) with a random duration that was uniformly sampled from 30 to 45 seconds.

Second, in the 20 minutes training phase, both the update of visual patterns and the delivery of electroshocks were dependent upon fish’s behavior. After the visual pattern was updated (including the first visual pattern in the training stage), fish had 7 seconds thinking-time to escape from the CS zone. If fish were in the CS zone after the thinking time, whole-arena shocks would be delivered every 3 seconds until fish escaped from the CS zone. After fish stayed in the Non-CS zone for 48 seconds, the visual patterns (CS zone at the top or bottom) would update with equal probability. The whole procedure would repeat (Figure 1B).

After the training phase, there was a one-minute blackout phase to deprive all visual stimuli.

Finally, in the last 18 minutes test phase, to ask whether fish could develop the association between the CS pattern and the US shock, the visual pattern interchanges every two minutes between at the top and at the bottom until the end.

#### 2.4.2 Self-control conditioning protocol

All phases were identical to the operant learning protocol, except for no electroshock delivery.

#### 2.4.3 Unpaired operant learning protocol

All phases were identical to the operant learning protocol except for the training phase, in which electroshocks, without pairing with visual patterns, were randomly delivered across the 20-minute duration.

### 2.5 Behavioral Analysis

#### 2.5.1 Visual intensity ratio

The visual intensity ratio is defined as the ratio of the grayscale value of the conditioned pattern to the grayscale value of the pure-gray pattern (the non-conditioned pattern). The descending ranking of intensity ratios: the white-black checkerboard > the red-black checkerboard > the pure-black pattern (see Table. 1 for more details).

**Table 1.** Visual intensity ratios of all visual patterns to the pure-gray pattern. In the table, each column stands for a visual pattern used in the experiment and rows show the mean values, the grayscale values, and the contrast ratios to the pure-gray pattern (the pattern at the non-CS zone, see Methods).

|  | White-black checkerboard | Red-black checkerboard | Pure black pattern | Pure gray pattern |
| --- | --- | --- | --- | --- |
| Mean RGB value | (128,128,128) | (128,0,0) | (0,0,0) | (128,128,128) |
| Grayscale value | 128 | 43 | 0 | 128 |
| Intensity ratio to the pure-gray pattern | 1 | 0.34 | 0.0 | 1 |

#### 2.5.2 Pre-screening

We define data quality as the percentage of not-bad frames. Frames when fish froze over 1 second were considered bad. Fish with data quality lower than 0.95 were excluded from the analysis since those fish did not swim spontaneously and frequently. Those fish were considered not in good conditions.

The positional index is defined as the percentage of frames when fish were in the non-CS zone.

#### 2.5.3 Turning analysis

We scored a turning event when the heading angle change between two consecutive frames exceeded 15 degrees. The fish would get +1 score when performing an escape turn, and −1 score when returning to the CS zone. Fish in the Non-CS zone executed an escape turn when they approached the midline (within twice body length) and then turned back (Figure S2). The turning index is defined as

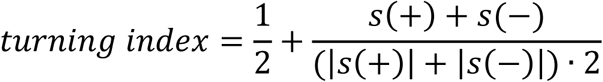

where s(+) and s(-) are positive and negative scores respectively. In this way, the turning index would fall between 0 and 1, the same range as the positional index.

#### 2.5.4 Distance to the mid-line

This is defined as a signed Euclidean distance from the fish head position to the mid-line. The sign is −1 when fish were in the CS zone and +1 when fish were in the non-CS zone.

#### 2.5.5 Learning analysis

To evaluate whether fish learned the operant learning task, we divided the entire operant conditioning protocol time into 24 two-minute-epochs. The memory may go extinct during the test phase in the absence of electroshocks (Myers and Davis, 2007). The extinction point was computed as the first time when the positional index within an epoch dropped below the baseline. The retrievable period was defined from the starting time of the test phase to the extinction point. We would use the memory length or the retrievable period interchangeably. If the positional indices in the retrievable period were significantly higher than the positional indices in the baseline phase, fish were classified as learners (The unpaired t-test was applied).

The positional index increment is the difference between the mean positional index in the retrieval period and the mean index in the baseline period. And the turning index increment is the difference between the mean turning index in the retrieval period and the mean index in the baseline period. The learning ratio is the ratio of the number of learners to the total number of fish.

### 2.6 Statistical Analysis

The paired t-tests were used to compare the difference between fish trained with the self-control conditioning protocol and the operant conditioning protocol; whereas the unpaired t-tests were used for the comparison between fish trained with the unpaired operant control protocol and those with operant learning protocol. The sample size exceeded 20 for all tests.

### 2.7 Linear Regression

The linear regression model

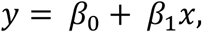

where *β*_0_, *β*_1_ were linear coefficients, was used to statistically quantify the trend of learning versus ages in terms of memory length, positional index increment, and turning index increment. Estimated linear coefficients, R-squared coefficients, and p-values for F-tests on the model were calculated using fitlm in the MATLAB Statistics and Machine Learning Toolbox.

## 3 Results

### 3.1 Larval zebrafish show significant learning responses in the operant learning task

#### 3.1.1 A high-throughput behavioral system for the operant learning task

In our modified operant learning task (Figure 1B), larval zebrafish freely swam in an arena divided by two distinct patterns, each of which was projected onto one half of a transparent floor. In all cases, a pure-gray visual pattern was presented in the non-CS zone, whereas other patterns were presented on the other half as the CS. The CS was paired with the US, moderate electroshocks. The delivery of the US and the update of visual patterns depended upon fish’s positions (see Material and Methods for detailed experimental procedures).

To scale up the learning process, we developed a high-throughput operant conditioning system (Figure 1A) with supporting software suites BLITZ and ABLITZER (see Material and Methods) that allowed training 12 fish simultaneously. BLITZ provided a fully automated workflow from video capture, online image processing, to visual stimulus presentation and electroshocks delivery for all behavioral protocols. Raw experimental data were then imported, analyzed and visualized by ABLITZER.

#### 3.1.2 Larval zebrafish show significant learning responses in the operant learning task

We found that 7-10 dpf zebrafish larvae showed significant learning responses (Figure 1C and Figure 1D), evaluated based on fish positions — positional index and turning index (see Material and Methods). Because larval zebrafish have the innate positive light preference, we developed two control settings: a self-control conditioning protocol in which no electroshock were delivered and an unpaired operant learning protocol in which electroshocks were randomly delivered (see Material and Methods). Results from the two control settings were compared with those from operant learning protocol to determine whether fish learned the association. Figure 1E shows a representative trajectory of a learner who tended to avoid conditioned visual pattern after training.

### 3.2 Visual intensity ratio modulates operant learning responses in larval zebrafish

We asked whether visual intensity ratios — CS patterns with different mean intensities to the non-CS pattern (pure-gray pattern) — would modulate learning. Indeed, the lower CS to non-CS intensity ratio led to stronger learning responses: the group of fish presented with the white-black checkerboard showed almost no learning response (Figure 2A and Figure 2B), whereas those presented with the pure-black pattern showed prominent learning responses (Figure 2C and Figure 2D), quantified by the positional index and turning index. The learning indices for fish presented with the red-black checkerboard fell in between the two other cases (Table 2).

**Table 2.** Comparison of learning response increment between different conditioned patterns. In the table, each column stands for each conditioned visual stimulus used in the experiment and rows show the positional index increment and the turning index increment. (t-test)

|  | White-black Checkerboard | Red-black Checkerboard | Pure Black Pattern |
| --- | --- | --- | --- |
| Positional Index Increment | 0.0015 (p = 0.8832) | 0.0386 (p = 0.0021) | 0.1010 (p = 5.92 x 10<sup>-6</sup>) |
| Turning Index Increment | -0.0239 (p = 0.3705) | 0.0599 (p = 0.0099) | 0.1103 (p = 0.0057) |

### 3.3 Single fish analysis distinguishes learners from non-learners

After the population analysis of learning responses, we next focused on individuals, e.g., to count learners. Here, we divided the entire process into epochs. Every two-minute-interval was one epoch. Therefore, the baseline phase has five epochs; the training phase has ten epochs, and the test phase has nine epochs.

#### 3.3.1 Memory extinction

We define the memory extinction point as the first time when the positional index within an epoch drops below the index in the baseline phase, and define the duration from the start of the test phase to the extinction point as the memory length. Memory length shorter than two epochs (e.g., fish may stay still in the non-CS zone) were excluded (see Material and Methods).

#### 3.3.2 Single fish analysis

Fish were categorized as learners only when their positional indices across the memory length were significantly higher than the indices in the baseline phase (see Material and Methods). We found that 26% of the fish were learners when the CS was red-black checkerboard (N = 104), and 50% of the fish were learners when the CS was pure-black pattern (N = 42). When white-black checkerboard was used as the CS, only one out of 37 fish learned (Table 3). The learners’ group showed significant changes in both the mean positional index and turning index before and after training (Figure 3B, C and Figure 3E, F).

**Table 3.** Age-dependent learning ratio and memory length (CS zone was red-black checkerboard) In the table, each column stands for fish age and rows show the learning ratio and memory length.

|  | 7 | 8 | 9 | 10 | Total |
| --- | --- | --- | --- | --- | --- |
| Learning Ratio | 4/16 | 4/19 | 6/29 | 13/40 | 27/104 |
| Memory Length (s) | 450 | 690 | 740 | 877 | 756 |

**Table 4.**
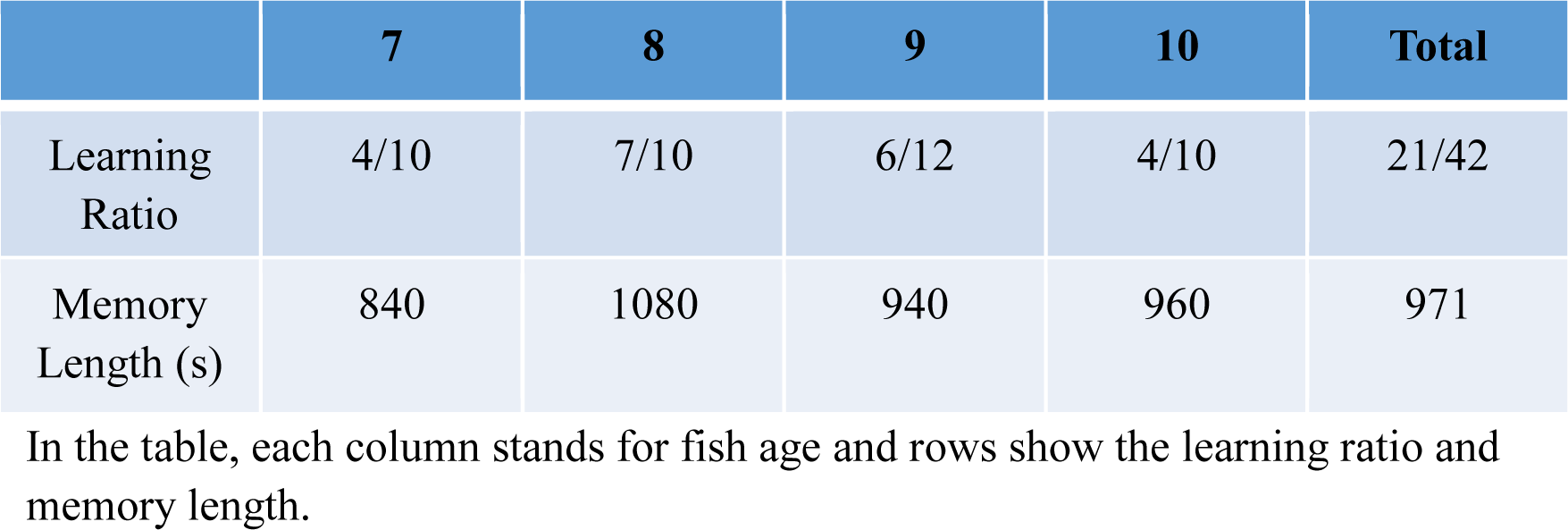
Age-dependent learning ratio and memory length (CS zone was pure-black pattern) In the table, each column stands for fish age and rows show the learning ratio and memory length.

|  | 7 | 8 | 9 | 10 | Total |
| --- | --- | --- | --- | --- | --- |
| Learning Ratio | 4/10 | 7/10 | 6/12 | 4/10 | 21/42 |
| Memory Length (s) | 840 | 1080 | 940 | 960 | 971 |

**Figure 3.**
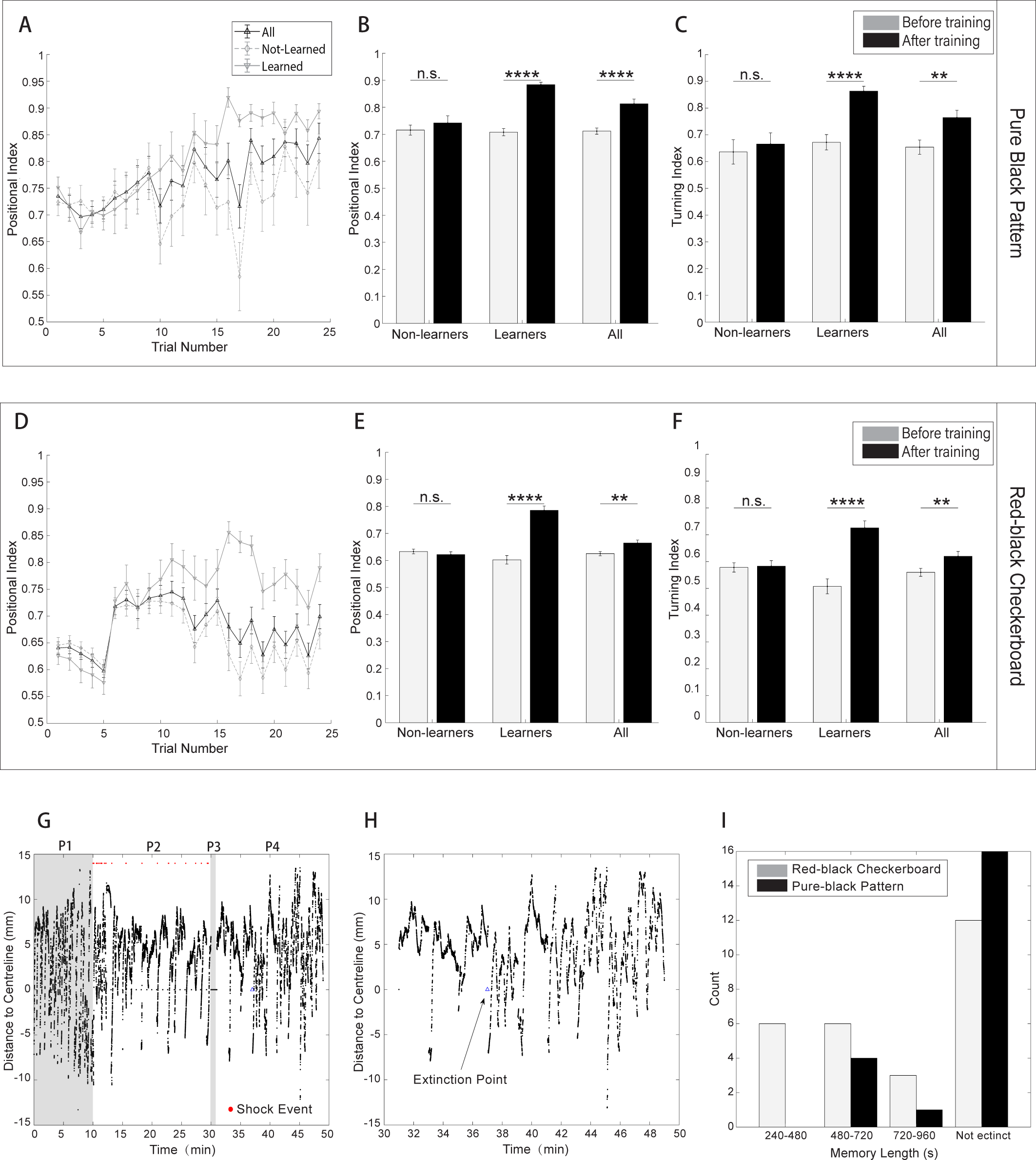
Single fish analysis distinguishes learners from non-learners. (A) Positional index averaged over all fish in the experiment group as well as the subpopulations classified as learners (light solid line, N = 21) and non-learners (light dash line, N = 21). CS zone was black pattern and the entire training process was divided into two-minute-epochs. (B) Analysis of the positional index suggested that the learners (N = 21) showed significant learning responses before and after training; whereas the non-learners (N = 21) did not show significant learning responses. (CS zone was black pattern; t-test, p = 1.98e-11 for the learners, p = 0.9492 for the non-learners and p = 2.03e-06 for all fish.) (C) Learners also showed significant difference in the turning indices. (CS zone was black pattern; t-test, p = 1.22e-5 for the learners, p =0.6491 for the non-learners and p = 0.0057 for all fish.) (D) Positional index averaged over all fish in the experiment group as well as the subpopulations classified as learners (light solid line, N = 27) and non-learners (light dash line, N = 77). CS zone was red-black checkerboard; same analysis as in (A). (E) Learners showed significant difference in the positional indices, the same analysis as in (B). (CS zone was red-black checkerboard; t-test, p = 6.52e-12 for the learners, p =0.3849 for the non-learners and p = 0.0020 for all fish.) (F) Learners also showed significant difference in the turning indices, same analysis as in (C). (CS zone was red-black checkerboard; t-test, p = 2.28e-6 for the learners, p = 0.8606 for the non-learners and p = 0.0099 for all fish.) (G) A typical trace of a learner whose memory would extinct during the test phase (CS zone was red-black checkerboard). The blue triangle denotes the extinction point when the single-epoch-averaged positional index drops below the mean index of the baseline. (H) A magnification of the test phase in (G). (I) Distributions of memory lengths of learners when CS zone was red-black checkerboard or pure black pattern respectively.

We plotted the learning curve — positional indices versus time — for learners and non-learners (Figure 3A). In the case of red-black checkerboard learners, the learning curve rose and approached the maximum near the end of training; during the test phase, the learning curve remained high during the first three epochs before memory extinction. In Figure 3G, we showed the trace of a typical fish with memory extinction in the test phase. Figure 3H magnified the test phase of Figure 3G, in which after the extinction point at ∼ 37 minute, fish started to swim more in the CS-zone.

In the case of pure black pattern learners, the learning curve also reached its maximum near the end of training. However, it remained high across the entire test phase (Figure 3D).

In Figure 3I, we compared the distribution of memory lengths when two different CS patterns were used. The mean memory length in the red-black checkerboard case was 756 seconds whereas the mean memory length lasted 970 seconds in the pure-black pattern case. Also, all black pattern learners’ memory lengths were longer than 480 seconds.

### 3.4 Age-dependent operant learning in larval zebrafish

We evaluated the learning abilities across 7-10 dpf larval zebrafish by plotting the memory length, positional index and turning index versus ages.

In the case of learning red-black checkerboard pattern, the positional index increment (see Material and Methods) and the memory length shows an increasing trend from 7 dpf to 10 dpf (Figure 4A and Figure 4B). However, there is no such trend in the turning index increment (see Material and Methods and Figure 4C).

**Figure 4.**
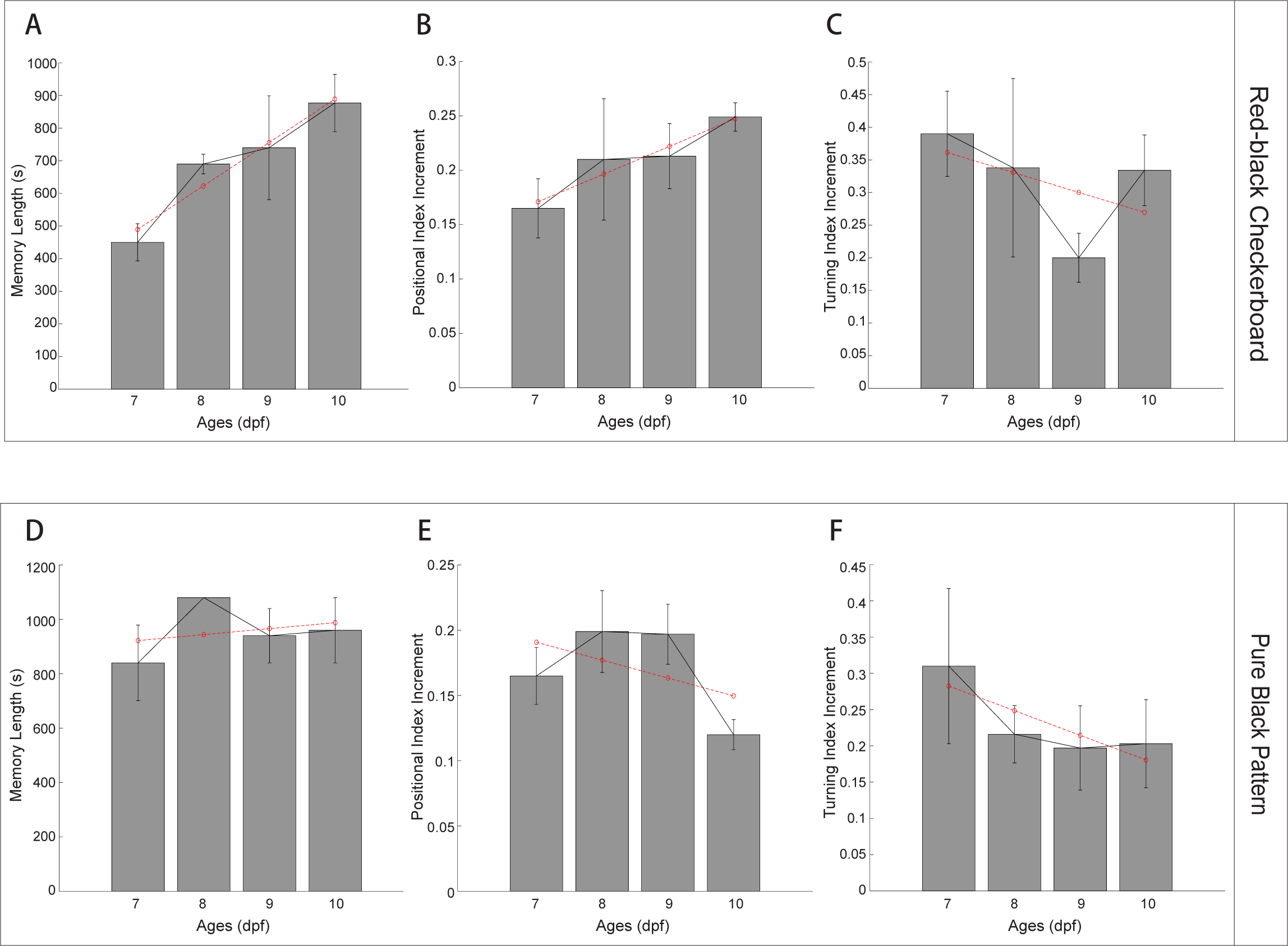
Age-dependent operant learning ability in larval zebrafish. (A) The mean memory length of all learners at specific age (CS zone was red-black checkerboard). Error bars are S.E.M. Linear regression was applied (red dashed line) to show the statistical trend. (Bi-square fitting applied, R-square = 0.932, p = 0.0347) (B) The mean positional index increment (CS zone was red-black checkerboard), the same analysis as in (A). (Bi-square fitting applied, R-square = 0.915, p = 0.0434) (C) The mean turning index increment (CS zone was red-black checkerboard), the same analysis as in (A). (Bi-square fitting applied, R-square = 0.237, p = 0.5130) (D) The mean memory length of all learners at specific ages (CS zone was pure-black pattern), same analysis as in (A). (Bi-square fitting applied, R-square = 0.033, p = 0.8190) (E) The mean positional index increment (CS zone was pure-black pattern), the same analysis as in (A). (Bi-square fitting applied, R-square = 0.229, p = 0.5210) (F) The mean turning index increment (CS zone was pure-black pattern); the same analysis as in (A). (Bi-square fitting applied, R-square = 0.688, p = 0.1710)

In the case of learning the black visual pattern, however, no increasing trends from 7 dpf to 10 dpf fish were found for the memory length (Figure 4D), the positional index increment (Figure 4E) and the turning index increment (Figure 4F).

## 4. Discussion

### 4.1 Operant learning in larval zebrafish

Operant learning allows animals to avoid dangers or to find potential rewards in a complex environment (Skinner, 1984). In this study, we demonstrated 7-10 dpf larval zebrafish showed significant operant learning responses when the CS, for example a red-black checkerboard pattern, was paired with the US, noxious electroshocks. In an earlier study (Valente et al., 2012), it was reported that one-week larvae showed no significant learning response. Several factors may explain this discrepancy. First, we observed little learning response when the white-black checkerboard was paired with the US (only one fish learned the contingency), consistent with Valente’s results. Enhancement of learning was observed, however, when the red-black checkerboard was paired with the US. In both cases, the non-CS zone was pure gray. The red-black checkerboard has a lower visual intensity ratio than the white-black checkerboard (see Table 1), and we hypothesize that visual intensity could strongly modulate the learning response. Second, in our modified paradigm, fish had more opportunity to learn the contingency between the CS and the US during the training period: when fish stayed in the non-CS zone for more than 48 seconds, the positions of the CS and non-CS patterns would update. In Valente’s paradigm, however, there were no visual pattern updates when fish stayed within the non-CS zone.

### 4.2 Visual intensity ratio modulates learning in larval zebrafish

We further investigated whether different visual intensity ratios could modulate learning ability in larval zebrafish. We found that fish showed little learning response when the white-black checkerboard was used as the CS pattern, which has the same average intensity as pure gray, the non-CS visual pattern. However, when the red-black checkerboard or pure-black visual pattern was used, some fish (28% in the group exposed to the red-black checkerboard and 50% exposed to the pure-black pattern) showed strong learning responses. Moreover, both the positional index and turning index were significantly higher in fish exposed to pure-black visual pattern versus those exposed to the red-black checkerboard (see Table 2).

Many studies have demonstrated that larval zebrafish exhibit positive phototaxis (Steenbergen et al.2011; Chen and Engert 2014; Guggiana-Nilo and Engert 2016). In our behavioral paradigm, the behavioral metric baselines (e.g., positional index) were computed first from a self-control procedure (see Material and Methods) before the operant learning procedure started. Visual intensity ratio could shift the baselines (see Table 2) due to animal’s innate bias. Significant changes of the behavioral metrics during and after operant learning (see Figure 2), however, require an explanation that goes beyond innate avoidance responses.

Here we speculate that intensity-ratio-dependent learning may arise from the crosstalk between the phototaxis and fear learning circuits. Both the phototaxis and US-triggered fear responses involve habenula (Agetsuma et al., 2010; Zhang et al., 2017), a specialized brain region where a direct association of the CS with fear may occur through synaptic plasticity. According to this model, the CS would trigger fear responses, and learning leads to a stronger association of the visual-related input and the escape response. These predictions can potentially be tested by combining our behavioral system with whole brain calcium imaging in freely behaving larval zebrafish (Cong et al., 2017).

### 4.3 Memory extinction

Memory extinction is an active learning process where an animal learns to dissociate the conditioned response and the CS in the absence of the US (Myers and Davis, 2007). In our assay, the extinction point is defined as the first epoch whose positional index dropped below the mean positional index of the baseline. In addition, fish that did not keep a high level of the positional index for at least two epochs were not counted (see Material and Methods). When the red-black checkerboard was used as the CS, memory lengths were distributed within a range of 7-18 minutes (Figure 3I), consistent with a recent classical conditioning paradigm in larval zebrafish (Aizenberg and Schuman, 2011). When the pure-black pattern was used as the CS, few learners showed memory extinction before the 18-minute test phase ended (see Figure 3G). Taken together, these results suggest that both operant conditioning and memory extinction could be differentially modulated by the visual intensity ratio.

### 4.4 Development of Operant Learning in Larvae

We selected 7-10 dpf zebrafish larvae for operant conditioning, a choice that was based on two considerations. First, larvae at 6 dpf show frequent long-pauses (over 7 seconds), and therefore are not suitable for operant conditioning: an animal must explore the action space instead of staying still. Second, an earlier work (Ingebretson and Masino, 2013) found that larvae at 7 dpf and later will produce more consistent locomotor activities. Here we found that 10 dpf fish exhibited the longest memory length when the red-black checkerboard was used as the CS; whereas 7 dpf fish showed the shortest memory length when the pure-black pattern was used (see Figure 4A). In addition, in the case of associating the red-black checkerboard pattern with the US, there is an age-dependent increasing trend for the memory length (Figure 4A) and the positional index increment (Figure 4B). No such trends were found when the pure-black pattern was used as CS (see Figure 4D, E, and F). These differences may partially result from a continuous development of larval zebrafish brain (Mueller and Wullimann, 2013).

### 4.5 High-throughput behavioral assays for learning and memory in Larval Zebrafish

Larval zebrafish are amenable to high-throughput screen due to their transparency, small size and high permeability to small molecules (Kokel et al., 2010; Rihel et al., 2010). Though most high-throughput systems are designed for drug or genetic screens (Gehrig et al., 2018; Rihel et al., 2010; Yang et al., 2018), here we have developed a high-throughput behavioral training system with custom supported software suites. Compared with previous work (Hinz et al., 2013; Pelkowski et al., 2011), the BLITZ software has enabled a fully automatic control of video capture, online image processing, visual pattern presentation and electroshocks delivery, making it an easily adaptable system for various purposes. Our complementary ABLITZER software also allows users to import, analyze and visualize data with well-structured classes and functions.

So far, our system cannot deal with situations of overlapping larvae, whose identities are hard to assign based on the current tracking algorithm. An earlier work (Mirat et al., 2013) showed that accurately tracking multiple larvae in groups over long periods of time were feasible. Integration of their algorithm with BLITZ may allow the study of social interactions of larval zebrafish in the future (Buske and Gerlai, 2014).

In conclusion, we have developed a high-throughput operant conditioning system for larval zebrafish. When using electroshocks as the US and red-black checkerboard or pure-black pattern as the CS, we demonstrated that a fraction of larval zebrafish could acquire operant learning, and the performances strongly depended on the visual intensity ratio of the CS to the non-CS pattern. Finally, we also identified age-dependent learning variability across 7-10 dpf larval zebrafish.

## 6 Author Contributions

Wenbin Yang conceived the study, designed and built the behavioral setup, developed the software suites, designed, carried out the experiments, wrote the manuscript and conceived the figures.

Yutong Meng helped carry out the experiment and conceive the figures. Danyang Li helped design, build the behavioral setup and conceive the figures. Quan Wen helped design the experiments and wrote the manuscript.

## 7 Funding

This work was funded by National Science Foundation of China Grants NSFC-31471051 and NSFC-91632102, the Strategic Priority Research Program of the Chinese Academy of Sciences (Pilot study, grant XDPB10).

### 8 Acknowledgments

The authors would like to thank Bing Hu for kindly providing the housing for zebrafish, Yuming Chai, Kexin Qi and Kun He for invaluable technical help; Florian Engert, Caroline Wee, Max Nikitchinko and Armin Bahl for kind guidance and inspirations in behavioral neuroscience.

## 9 Supplementary Material

The Supplementary Material for this article can be found in the attachment.

## 10 Abbreviations

dpf: days post fertilization
fps: frames per second
SEM: standard error of the mean
BLITZ: behavioral learning in the zebrafish
ABLITZER: the analyzer of BLITZ results

## 11 Data Availability Statement

Datasets are available on request.

